# Presence of polyadenylated 3’ tail in RNA of *Pepper mild mottle virus* allows the Oligo(dT)_18_ in Priming cDNA Synthesis of its genomic RNA template

**DOI:** 10.1101/2020.04.13.039180

**Authors:** Nidhi Kumari, Vivek Sharma, Sneha Choudhary, P.N. Sharma

## Abstract

The PCR amplification of majority of the ssRNA of both genomic and non-genomic mRNA is accomplished by RT-PCR. The mRNA is subjected to cDNA synthesis using reverse transcriptase and either Oligo(dT)18, or random or gene specific reverse primers based priming strategies. The choice of primer largely depends on the nature of 3 prime terminus of mRNA and length of cDNA synthesized. In general, oligo(dT)18 is the preferred choice for mRNAs having poly(A) tail at 3 prime terminus. In general, tobamoviruses lack any poly(A) tail at their 3 prime untranslated region (UTR) which forms a tRNA like structure and upstream pseudoknot domain except tow viruses viz., Hibiscus latent Fort Pierce virus (HLFPV) and Hibiscus latent Singapore virus (HLSV) which accommodate internal poly(A) sequences of 46 and 87 nucleotides long, respectively in their 3 prime UTR. However, determination of full nucleotide sequence of Pepper mild mottle virus (PMMoV) using an oligo(dT)18 primed cDNA as template indicated the libertinism of oligo(dT)18 in priming cDNA synthesis of RNA template which are known to lack poly(A) tail. at the end or internally at its 3 prime end. Moreover, coat protein (CP) gene of PMMoV and bean common mosaic virus (BCMV) (Potyvirus with a poly(A) tract at its 3 prime end) was amplified using cDNA primed with random primer as well as oligo(dT)18 was successfully amplified but with significant variation in the intensity of the amplification band in case of PMMoV but not in BCMV. This clearly indicated the presence of PMMoV mRNA with polyadenylated 3 prime tail in total RNA isolated from PMMoV infected capsicum leaves with abundance of non-polyadenylated PMMoV genomic RNA (gRNA). Hence, we hypothesize that the generation of polyadenylated RNA population during the infection cycle of PMMoV in pepper may be possible reason for libertinism of oligo(dT)18 in priming cDNA synthesis of RNA template isolated from PMMoV infected leaves followed by amplification of entire PMMoV genome through RT-PCR. This is first study indicating the presence of polyadenylated or polyadenylated rich regions in PMMoV gRNA acquired during the infection cycle in pepper.

## Introduction

Plant viruses cause significant deviation from normal growth and development in plants leading to heavy losses in the quality and quantity of plant produce and therefore hold an economic importance (Gergerich and Dolja, 2006). Out of nearly 1000 described plant viruses, about 800 contain single stranded (ss) RNA as their genetic material (Hull, 2002). The positive sense ssRNA represents the largest class of the plant viruses (Ahlquist, 2002). The RNA viruses contain a variety of structures both at 5’ and 3’ ends of their genomes. At 5’ terminus, plant viruses may have phosphate group, a cap or a virus-encoded polypeptide called VPg (Viral Protein genome-linked) covalently attached to the first nucleotide of the RNA (Thivierge et al., 2005). The 3’ end of the RNA either features a tRNA-like structure (TLS) or a poly (A) tail or simply a 3’ hydroxyl (OH) group (Thivierge et al., 2005; Dreher, 1999). RNA polyadenylation involving homo/hetero polymeric poly (A) addition is a common feature of eukaryotic mRNA and some viral mRNAs in which length can vary from 20-100 nucleotides (Hull, 2002; Li et al., 2014). The post-transcriptional adenylation of homo or hetero poly(A) residues plays self-contradictory role in protecting transcripts from degradation, nucleocytoplasmic export and translation initiation (Moore and Proudfoot, 2009). Plant viruses which are reported to possess 3’ poly(A) tail includes *Potexviruses*, *Mandariviruses*, *Allexiviruses*, *Capilloviruses*, *Carlaviruses*, *Citriviruses*, *Foveaviruses*, *Tepoviruses*, *Trichoviruses*, *Vitiviruses*, *Marfiviruses*, *Maculaviruses*, *Potyviruses*, *Ipomoviruses*, *Macluraviruses*, *Poaceaeviruses*, *Rymoviruses*, *Tritimoviruses*, *Bymoviruses*, *Benyviruses*, *Cileviruses* (King et al., 2012). However, recently poly(A) tails at 3’ terminus of some viral RNA has also obtained in case of viruses which are not reported to bear a poly(A) tails (Li et al., 2014; He et al., 2015).

For genome amplification of RNA virus, the primary step is the cDNA synthesis. The RNA priming strategy depends on the nature of the 3’ terminus of mRNA and also plays an important role in determining the efficacy and length of cDNA synthesized. In general, three types of primers viz. oligo(dT)_18_, random primers or a sequence-specific reverse primer are used for cDNA synthesis. Each primer offers its own benefits as well as limitations. The most preferred method in case of viral mRNA bearing poly(A) tail at 3’ end is oligo(dT)_18_ priming, (Weiss et al., 1976; Verma, 1978) however, it may often leads to the formation of truncated cDNA in high frequency due to the internal poly(A) priming which could be minimized by simple replacement of the oligo(dT)_18_ primer with a set of anchored oligo(dT) primers (5’-ACTATCTAGAGCGGCCGCTTT16-A, -G, -CA, -CG, and -CC-3’) for cDNA synthesis (Nam et al., 2002).

The complete genome of PMMoV, has been determined using oligo(dT)_18_ primed cDNA in our previous study (Rialch et al., 2015). Being a tobamovirus, the 3’ UTR of PMMoV genomic RNA adopt a TLS, thus lack a poly(A) tail at 3’ terminus (Dreher, 1999; Rietveld et al., 1984; Belkum et al., 1985). In case of some other plant viruses without 3’ poly (A) tail also, oligo(dT)_18_ has successfully synthesized the cDNA from their genomic RNA (gRNA) (He et al., 2015, Li et al., 2014; Pyle et al., 2019). This raises the query that how oligo(dT)_18_ can prime cDNA synthesis of PMMoV, a tobamovirus that lacks 3’ poly(A) tail. Therefore, the present study was undertaken using two RNA plant viruses one belonging to genus *Potyvirus* with a known poly(A) tail (BCMV) and the other from *Tobamovirus* genus, characterized by the absence of any poly(A) tail at its 3’ terminus (PMMoV) to comprehend the libertinism of oligo(dT)_18_ in cDNA synthesis from PMMoV gRNA.

## Materials and Methods

### Total RNA isolation, 3’ RACE PCR and sequencing

Total RNA was extracted from BCMV and PMMoV infected beans and capsicum leaves, respectively using Trizol reagent following the manufacturer’s instructions. To determine the nature of 3’ end of both viruses, 3’-RACE-PCR was performed as per the instructions of SMARTer™ RACE-cDNA Amplification Kit User Manual (Clontech, 2012) using primers given in Table 1. For cDNA synthesis, initially in PCR tubes 1 μg total RNA (5 μl) and 3’ CDS primer were taken in total reaction of 10 μl, final volume was made by RNase free water, tubes were vortexed, centrifuged briefly to collect the mixture at the bottom and incubated at 70°C for 5 min followed by 2 min incubation on ice. After incubation, 4 μl of 5x MMLV buffer, 2 μl of 10 mM each dNTPs mix (Fermentas, USA), 1 μl of RNAase inhibitor (USB) and 1 μl (400U/μl) of MMLV reverse transcriptase were added to the reaction mixture and final reaction volume was made to 20 μl by adding nuclease free water. The contents were then incubated at 42°C for 90 min (Gen Amp PCR System 9700, Applied Biosystem, USA). PCR amplification was performed in 50 μl reaction volume using 5 μl of 10x Taq buffer, 3 μl of 25 mM MgCl_2_, 4 μl of 2 mM dNTPs mix, 5 μl of 10 mM UPM mix, 1 μl of gene specific primer (GSP) (Table 1), 2.5 μl of cDNA, 0.2 μl of 5U/μl Taq polymerase (Merck Genei) and final volume was adjusted to 50 μl with nuclease free water. Amplification was performed in GeneAmp PCR system 9700 (Applied Biosystems) with initial denaturation at 94°C followed by 5 cycles of 94°C for 30 sec and 72°C for 2 min, 5 cycles of 94°C for 30 sec, 68°C for 30 sec and 72°C for 2 min and final 27 cycles at 94°C for 30 sec, 60°C for 30 sec and 72°C for 2 min with final extension of 7 min at 72°C. The amplified fragment was purified, ligated into pGEMT-Easy vector and transformed into *E. coli* strain *DH5α*. The recombinant plasmid was isolated, lyophilized, custom sequenced (Xcelris Labs Ltd) and analyzed using NCBI – BLASTn analysis tool.

**Table 1:**
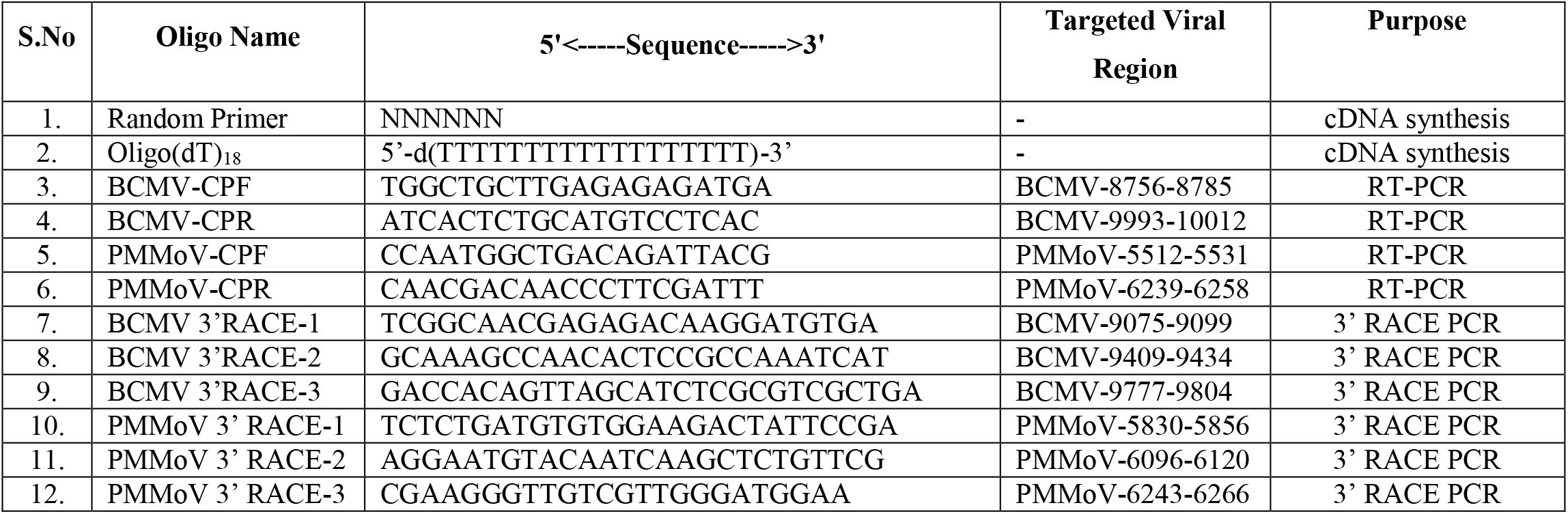
List of primers used for priming cDNA synthesis, RACE-PCR and RT-PCR

### cDNA synthesis and amplification of coat protein gene of PMMoV and BCMV

For cDNA synthesis, RNA was subjected to reverse transcription using random primer (NNNNNN) and Oligo(dT)_18_ (5’-TTTTTTTTTTTTTTTTTT-3’) as RNA reverse primer with Verso cDNA synthesis Kit as per the instructions of manufacturer at 42°C for 60 min followed by heat inactivation of reverse transcriptase at 95°C for 2 min. The cDNA synthesis of both the viruses was confirmed by RT-PCR using the CP specific primers of respective viruses (Rialch et al., 2015; Sharma et al., 2011). The RT-PCR was carried out as described by Rialch et al., 2015 and Sharma et al., 2011.

## Results and discussion

The RACE-PCR followed by nucleotide sequencing revealed the presence of 198 nt long 3’ UTR bearing a single A residue at 3’ terminus as last nucleotide of PMMoV (KJ631123) whereas in BCMV a poly(A) tail of 30 nucleotides was observed downstream the 218 nt long 3’ UTR (KY057338.1) (Fig. 1). The 3’ UTR obtained in present study matched with the reported PMMoV full nucleotide sequences determined by other workers from different parts of the world available at NCBI databases {AB069853.1 (Hagiwara et al., 2002);, AB550911 (Oliveira et al., 2010); AB000709 (Kirita et al., 1997), AJ308228.1 (Velasco et al., 2002); AB113117.1, AB113116.1 (Ichiki et al., 2005); AB126003.1 (Yoon et al., 2005); AY859497.1 (Wang et al., 2006); AB276030 (Genda et al., 2007). Like other tomaboviruses, the 3’UTR of PMMoV forms a t-RNA like structure along with an upstream pseudoknot domain (Guo and Wong, 2019). Though in case of tobamoviruses there are instances where the 3’ UTR of *Hibiscus latent Fort Pierce virus* (HLFPV) and *Hibiscus latent Singapore virus* (HLSV) accommodate internal poly(A) sequences of 46 and 87 nucleotides long, respectively (Yoshida et al., 2014; Srinivasan et al., 2005). However, in case of PMMoV, such internal poly(A) tracts were absent in case of all the 39 full genome sequences including the one determined in our previous study PMMoV-HP1 (KJ631123.1). Additionally, using the cDNA primed with random primer as well as oligodT(18) as template, CP gene of both the viruses was amplified. The CP gene of PMMoV and BCMV are up-stream of 3’UTR at 5685-6158 nt and 8876-9736 nt, respectively. The amplification product of ~730 bp and 1500 bp specific to the CP gene of respective viruses were obtained (Fig. 2). The successful amplification of CP gene with oligodT(18) primed cDNA clearly indicated the presence of PMMoV mRNA with polyadenylated 3’ tail in total RNA isolated from PMMoV infected capsicum leaves. Though there was considerable difference in the intensity of band in case of PMMoV with very thick and intense band of RT-PCR with random primer primed cDNA. However, band of same intensity was observed in case of BCMV with both random primer and oligo(dT)_18_ primers. This suggest that there is considerable variation in the population of non-polyadenylated and polyadenylated viral gRNA in total RNA isolated from PMMoV infected pepper leaves. Though it is evidential that the non-polyadenylated viral gRNA are more abundant. The possibility of synthesis of truncated cDNA has been ruled out as the complete PMMoV genome sequence was determined using the oligo(dT)_18_ primed cDNA as template. Previously, He et al. (2015) and Li et al. (2014) reported presence of poly(A) and poly(A)-rich tails in *Southern rice black-streaked dwarf virus* (SRBSDV) and *Tobacco mosaic virus* (TMV) belonging to genus *Fijivirus* and *Tobamovirus*, respectively. The RNA polyadenylation is a post-transcriptional modification of RNA which may have role in the RNA stability, guiding mRNA export to cytoplasm and even during translation initiation (Edmonds, 2002). The de-adenylation of a stable mRNA initiates its degradation by the action of exosomes or *hXrn1*. In contrast, transient poly (A) and poly (A) rich tails are considered as the degradation intermediates of RNA decay pathway that take place in both eukaryotic and prokaryotic cells (Slomovic et al., 2010; Regnier and Hajnsdorf, 2009; Lange et al., 2009). The widespread occurrence of poly (A) and poly (A) rich tails in case of virus genera lacking the 3’ poly (A) tails suggests their potential role in self-protection of viral gRNA from host mRNA degradation or surveillance mechanism (Pyle et al., 2019) or polyadenylation mediated viral RNA degradation as an antiviral defence mechanism of plant system (He et al., 2015). Therefore, we hypothesize that the generation of polyadenylated RNA population during the infection cycle of PMMoV in pepper was the basis for libertinism of oligo(dT)_18_ in priming cDNA synthesis of RNA template isolated from PMMoV infected leaves followed by amplification of entire PMMoV genome through RT-PCR.

**Fig 1:**
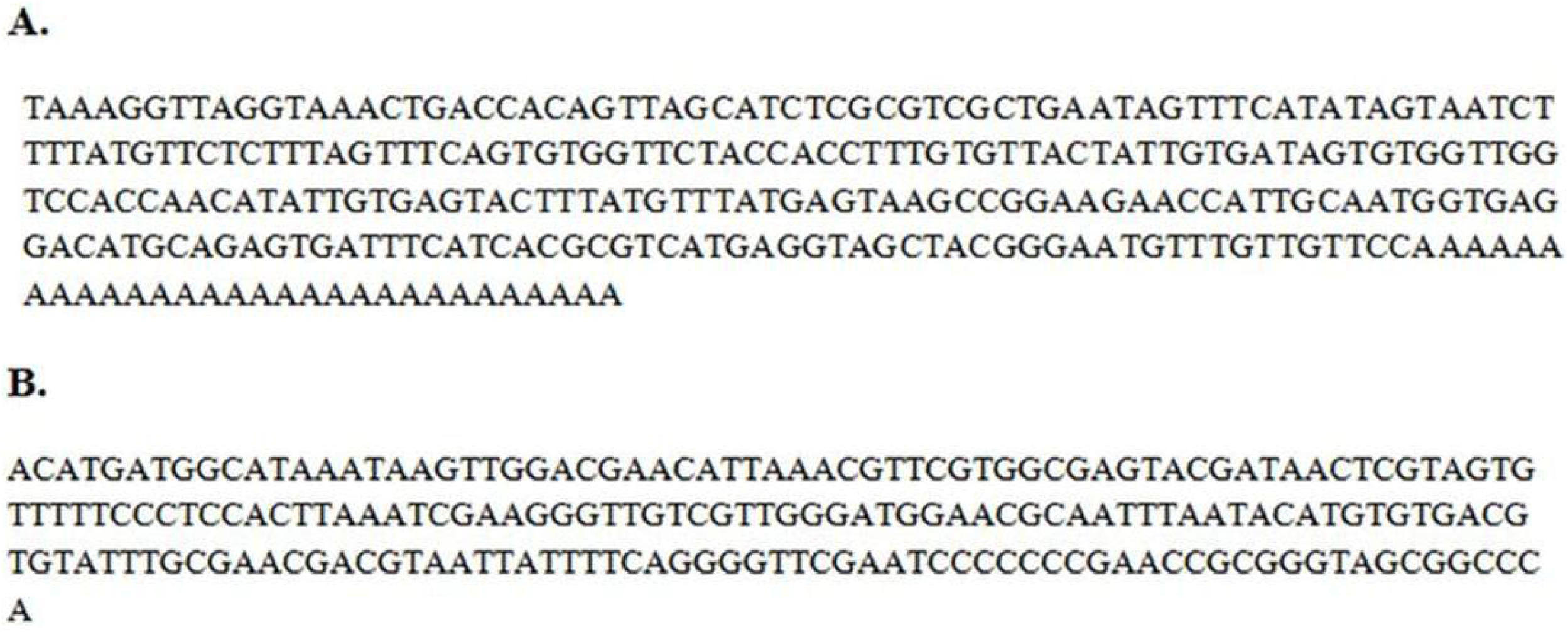
Nucleotide sequence of 3’ UTR of BCMV (A) and PMMoV (B) of 3’ RACE-PCR amplified products.

**Fig 2:**
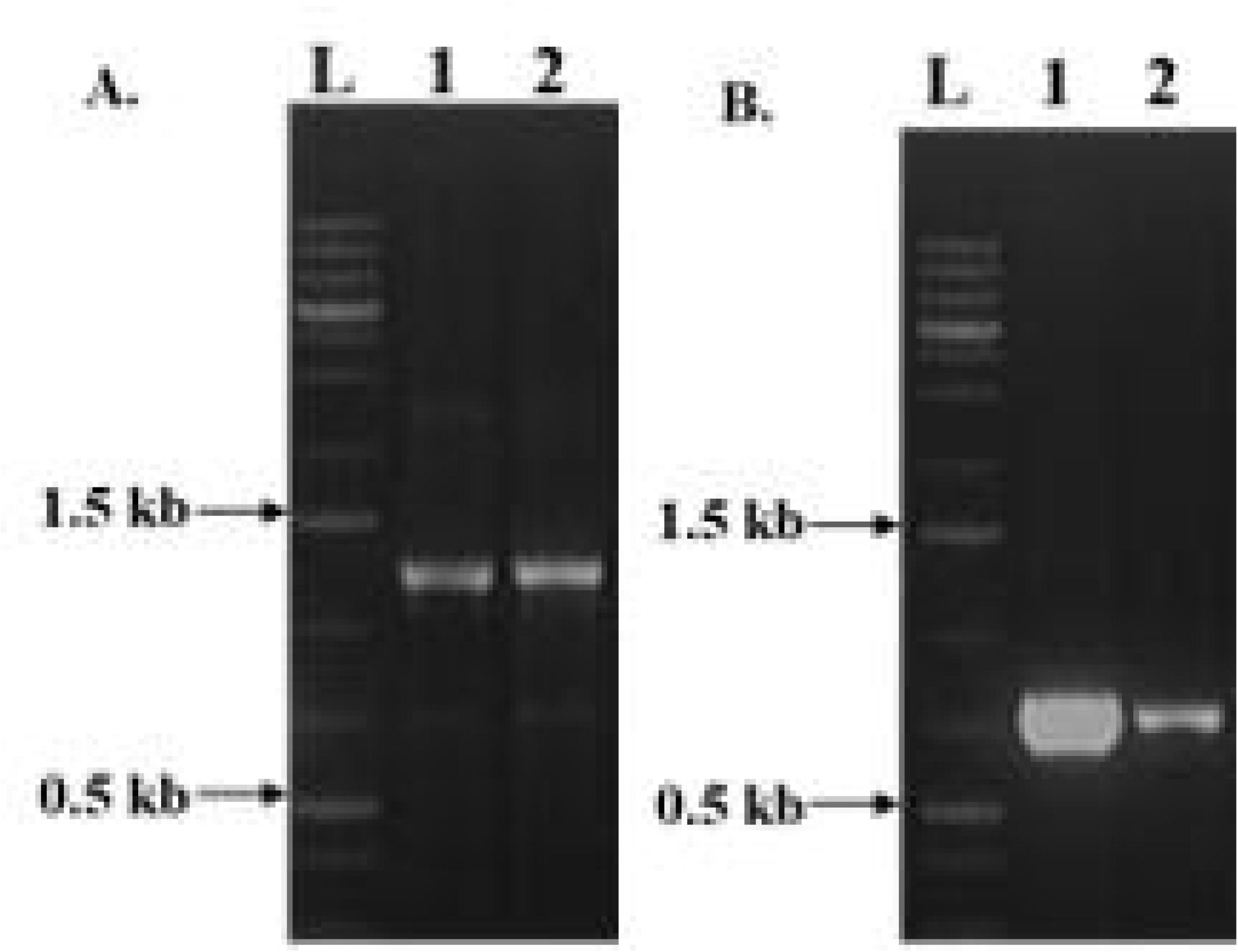
RT-PCR amplification of coat protein region of BCMV (A) and PMMoV (B) using cDNA synthesized from random primer (Lane 1) and oligodT_(18)_ (Lane 2).

## Acknowledgments

The senior author is thankful to University Grants Commission for providing the funds (award letter no.43-3/2014-SR) to carry out the present study.

## Compliance with ethical standards

**Conflict of interest:** None of the authors has any conflict of interest to declare.

